# Maximum Likelihood Estimation of Biological Relatedness from Low Coverage Sequencing Data

**DOI:** 10.1101/023374

**Authors:** Mikhail Lipatov, Komal Sanjeev, Rob Patro, Krishna R Veeramah

**Affiliations:** Department of Ecology and Evolution, Stony Brook University, Stony Brook, NY 11794; Department of Computer Science, Stony Brook University, Stony Brook, NY 11794

**Keywords:** 2^nd^ generation sequencing, low coverage, kinship, relatedness, SNPs

## Abstract

The inference of biological relatedness from DNA sequence data has a wide array of applications, such as in the study of human disease, anthropology and ecology. One of the most common analytical frameworks for performing this inference is to genotype individuals for large numbers of independent genomewide markers and use population allele frequencies to infer the probability of identity-by-descent (IBD) given observed genotypes. Current implementations of this class of methods assume genotypes are known without error. However, with the advent of 2^nd^ generation sequencing data there are now an increasing number of situations where the confidence attached to a particular genotype may be poor because of low coverage. Such scenarios may lead to biased estimates of the kinship coefficient, *ε* We describe an approach that utilizes genotype likelihoods rather than a single observed best genotype to estimate *ϕ* and demonstrate that we can accurately infer relatedness in both simulated and real 2^nd^ generation sequencing data from a wide variety of human populations down to at least the third degree when coverage is as low as 2x for both individuals, while other commonly used methods such as PLINK exhibit large biases in such situations. In addition the method appears to be robust when the assumed population allele frequencies are diverged from the true frequencies for realistic levels of genetic drift. This approach has been implemented in the C++ software *lcMLkin*.

## 2 Introduction

Biological relatedness can be quantified by a kinship coefficient (also known as the coancestry coefficient) [14], *ϕ*, that essentially quantifies the number of generations that separate a pair of individuals. More strictly, *ϕ* is the probability that two random alleles each selected from one in a pair of individuals are identical by descent (IBD). For example, parent-offspring and sibling-sibling pairs should possess *ϕ* of 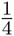, while for first cousins the value is expected to be 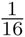,. Being the observed result of multi-generation geneaological process in a population, the extent of DNA sequence differences between two individuals is ideal data for inferring relatedness without any prior knowledge of the underlying pedigree, or when such knowledge is uncertain (see reviews [42, 38]). Such data is commonly used in a diverse array of fields such as the identification of disease-causing loci [10], forensics[4], anthropology [41], archaeology [12], genealogy [17] and ecology [36]. The higher *ϕ*, the more DNA sequence two individuals should share that is IBD. In a diploid population assumed to be outbred *ϕ* can be related to IBD through 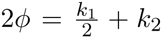, where *ϕ* is the coefficient of relatedness, *r*, and *k*1 and *k*2 are defined as the probabilities that two diploid individuals share 1 or 2 alleles that are identical by descent (IBD). In addition, one may also define *k*0 — the probability of two diploid individuals sharing 0 alleles that are IBD — such that *k*0 + *k*1 + *k*2 = 1. In the presence of inbreeding, additional *k* terms can be added [26, 42], though for sake of simplicity we ignore such scenarios. Thus, if the three *k* terms can be determined, it is possible to obtain an estimate of relatedness between two pairs of individuals.

Though the extent of IBD cannot be directly observed, it can be inferred from how much DNA is shown to be identical-by-state (IBS). The challenge, therefore, is to determine Pr(IBD*|*IBS). Other methods exist that model the transition of IBD along the genome [23, 18, 11, 6, 28, 8, 23], and the most common approaches use population allele frequencies to determine the likelihood of observing a particular genotype given a certain level of IBD at multiple loci and assume linkage equilibrium (i.e. independence) between individual sites [26, 38]. This framework has been applied to microsatellite loci with multiple alleles[15], and, with the advent of SNP microarrays, single base loci with two alleles (i.e. SNPs). The latter type of data, in particular, has power to infer relatedness down to at least fourth-degree relatives because of the large numbers of loci available. Method of moment estimators (e.g. PLINK [33], KING [25], REAP [39]) tend to be the most frequently used due to their ability to deal with large datasets at reasonable speeds (tens to hundreds of thousands of loci), though a maximum likelihood (ML) estimator was recently described for dealing with populations of mixed ancestry (RelateAdmix) [27].

In all of these current methods, genotype calls are assumed to be correct (or at least contain negligible error). However 2^nd^ generation sequencing is now emerging as the method of choice for obtaining genome-wide markers, either via whole genome shotgun or targeted capture. With 2^nd^ generation sequencing data, genotype quality is a function of sequencing coverage [13, 30]. While the ideal scenario is to obtain high genome coverage (*>*20X) for multiple individuals, this is not always feasible. Given a budget it may be preferable to sequence large numbers of individuals at low coverage, or samples may simply lack sufficient DNA material (for example in paleogenomic or forensic scenarios). Low coverage will lead to an underestimation of the true number heterozygotes, which may have the downstream affect of biasing subsequent estimates of the kinship coefficient.

In this paper, we describe a new method for inferring relatedness between pairs of individuals when the true genotypes are uncertain as a result of low-coverage 2^nd^ generation sequencing. Our approach is similar to other recent methods that attempt to infer population genetic parameters from low coverage data by utilizing genotype likelihoods rather than assuming a single best genotype [29, 20, 37, 19]. We show our method, implemented in the software *lcMLkin*, can accurately infer biological relatedness down to 5^th^ degree relatives from simulated data even when coverage is as low as 2x in both individuals examined. We then apply our method to real low-coverage 2^nd^ generation sequencing data and demonstrate that *lcMLkin* correctly estimates relatedness coefficients between individuals of known biological relatedness.

## 3 Materials and Methods

### 3.1 Model

Consider a single, non-inbred, non-admixed population for which there exist a biallelic locus with possible allelic states *B* and *C* and where the population allele frequencies are known. 2^nd^ generation sequencing data is generated at this locus (represented for example by an alignment of bases at this locus from sequence read data) for two individuals from this population with some degree of biological relatedness. Our goal is to use the sequence read data to estimate the relatedness coefficients for these individuals.

#### 3.1.1 Genotype Likelihoods

The (unknown) genotypes of the two individuals are designated by *G*^1^ and *G*^2^ and the three possible genotype values, *BB*, *BC* and *CC*, by *g*_0_, *g*_1_ and *g*_2_. The aligned sequence read data for individuals 1 and 2 at this locus are designated *N*^1^ and *N*^2^. The likelihood for each possible genotype for these two individuals given the read data can be expressed as:

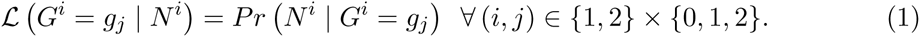

There are a number of different methods for calculating this likelihood that can account for factors such as independence or non-independence of reads, base and mapping quality or position of base call along the sequence read [24, 22, 9, 21, 16]. Unless stated, we use the method described by Depristo et al. [9], though ultimately this choice is up to the individual user.

#### 3.1.2 IBD/IBS Probabilities

Define *Z* as the number of alleles IBD between the two individuals at the biallelic locus — this is a latent variable in our model — and designate our estimate of the frequency of allelic states *B* and *C* in the source population by *p* and *q* = 1 –*p*. The probabilities of *Z* for 0, 1 or 2 given the observed pair of genotypes (i.e. given IBS) are well known [26]. Table 1 provides a relevant subset of these probabilities given the assumptions of our model (i.e. no inbreeding, no admixture, biallelic locus).

**Table 1:**
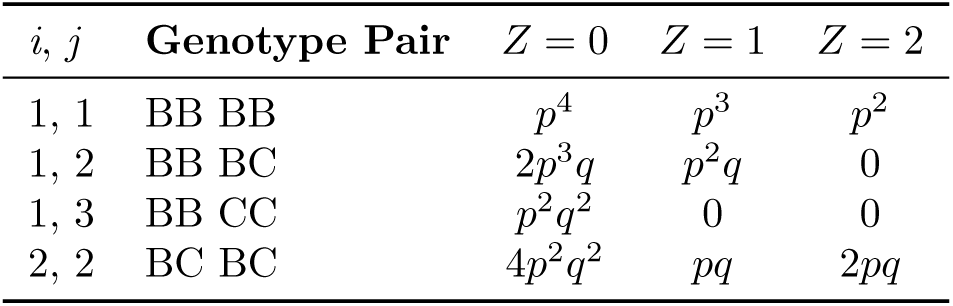
IBD probabilities for observed genotype pairs for individuals from the same unadmixed, non-inbred population

| $i, j$ | Genotype Pair | $Z = 0$ | $Z = 1$ | $Z = 2$ |
| --- | --- | --- | --- | --- |
| 1, 1 | BB BB | $p^4$ | $p^3$ | $p^2$ |
| 1, 2 | BB BC | $2p^3q$ | $p^2q$ | 0 |
| 1, 3 | BB CC | $p^2q^2$ | 0 | 0 |
| 2, 2 | BC BC | $4p^2q^2$ | $pq$ | $2pq$ |

The probability of a particular genotype combination does not change when we switch the individuals. Additionally, exchanging the identities of the two allelic states in a genotype combination amounts to exchanging *p* with *q* in the corresponding probability expression.

#### 3.1.3 Estimating the Kinship Coefficient

We define *K* as the 3-tuple of *k* coefficients, (*k*_0_, *k*_1_, *k*_2_). Note that 0 ≤ *k*_*z*_ ≤ 1 ∀*z ∈* {0, 1, 2} and that *k*_0_ + *k*_1_ + *k*_2_ = 1. Also note that *k*_*z*_ ≡ Pr (*Z* = *z* | *K*) ∀*z ϕ* {0, 1, 2}. We also define the combined kinship coefficient, 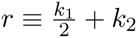.

Our approach for accounting for potential uncertainty in genotype calls because of low coverage 2^nd^ generation sequencing data when estimating *K* is to sum over all possible genotypes weighted by their likelihoods (i.e. we treat sequence reads as the observed data and genotypes as latent variables, which for the purposes of inference are effectively nuisance parameters) as in other recent methods attempting to estimate different parameters from such data. We can now write down a likelihood function for *K*, given *N*^1^, *N*^2^ and *p*for a given locus:

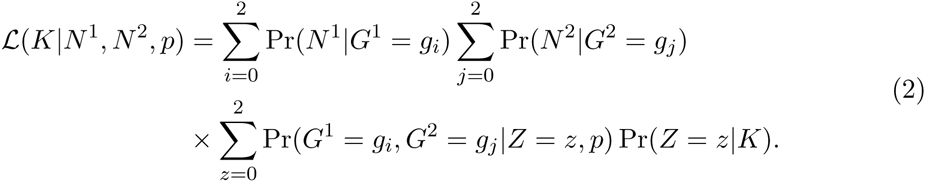

In our approach, we assume that all loci are in linkage equilibrium. Therefore, the total likelihood for a given *K* can be obtained from the product across loci (we take the sum of log likelihoods instead to avoid issues related to numerical precision). To obtain a maximum likelihood estimate of *K* (and thus also Фand *r*) we use an Expectation-Maximization (EM) algorithm. We also restrict the search space such that 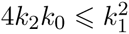.This method has been implemented in the C++ software *lcMLkin*(https://github.com/COMBINE-lab/maximum-likelihood-relatedness-estimation).

### 3.2 Data

#### 3.2.1 Simulated Pedigrees

Our aim was to simulate multiple pedigrees with the structure shown in Figure 1. This pedigree contains an array of relationships ranging from first degree 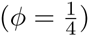 to fifth degree 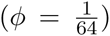 as well as unrelated or founder individuals. All population allele frequencies were obtained from samples genotyped at autosomal SNPs as part of the Human Origins Array [32]. To simulate a non-admixed population, allele frequencies were estimated from 100, 000 randomly chosen SNPs that were shown to have a minor allele frequency greater than 5% amongst 28 unrelated French individuals. Genotypes for each simulated locus for the 8 founders from each pedigree were binomially sampled given *p*. Genotypes in the other pedigree members were then sampled conditioned on these founder genotypes assuming Mendelian inheritance and independence between loci.

**Figure 1:**
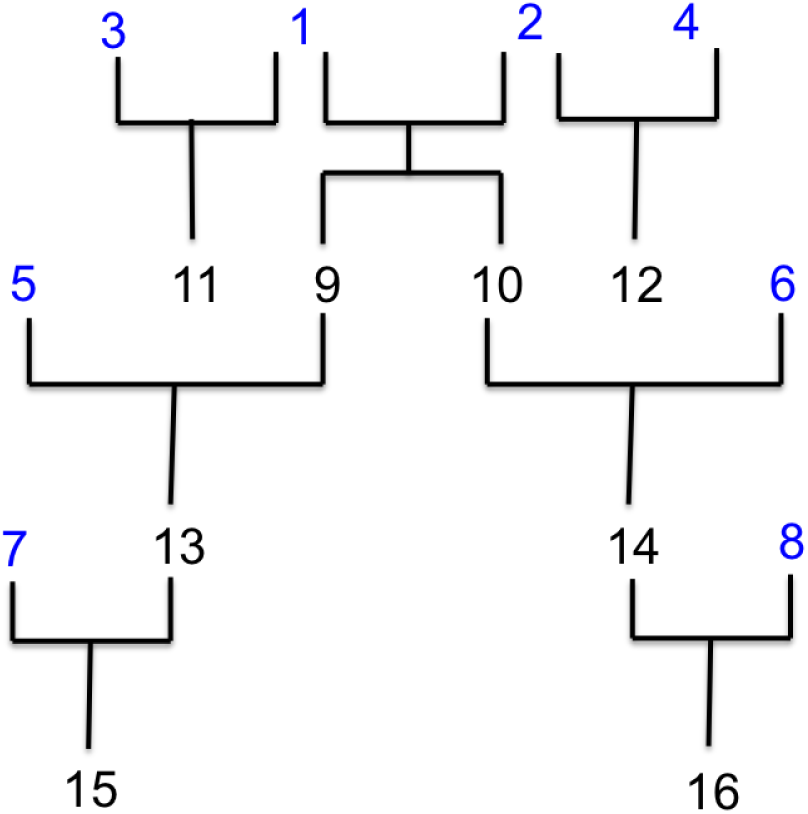
Topology of our simulated pedigrees. Individuals colored in blue are the unrelated founder individuals.

To simulate 2^nd^ generation sequencing data for an individual at a given mean coverage *x*, the number of reads for each locus is drawn from a poisson distribution with λ = *x*, and base calls for each read are randomly drawn given the individual’s true genotypes. Each base call is assigned a Phred quality score of 20 [35] and is changed to the opposite allele given this probability of an error (i.e. 1%). Thus, in our simulations, we only assume two possible alleles, rather than four. We experimented with more complicated quality score distributions, but found they did not change the results. We do not take into account mapping error for a read. A similar scheme is described in Veeramah et al. [40]. For each individual, we simulate 2^nd^ generation sequencing data at 2x-20x coverage in 2x intervals. Genotype likelihoods for each of the three possible genotypes are then calculated using the formula from Depristo et al. [9], accounting for the fact we only use two possible alleles.

#### 3.2.2 CEPH pedigree 1463

All 17 members of CEPH pedigree 1463 have been sequenced to high coverage (*∼* 50x) as part of Illumina’s Platinum Genomes dataset. BAM files were obtained for five of these individuals (NA12877, NA12883, NA12885, NA12889 and NA12890) such that there were pairs of known parent-offspring,sibling-sibling and grandparent-grandchildren relationships (http://www.ebi.ac.uk/ena/data/view/ERP001960). Approximately 10,000 SNPs were randomly selected from the Human Origins array subject to the requirement they were at least 250kb apart. For each of the five individuals, sequence reads at these SNPs were down-sampled into 10 new BAM files such that the mean coverage for each individual ranged from 2x-20x in 2x intervals. Genotype likelihoods for the three possible genotypes given the two alleles identified by the Human Origins array for each locus were then calculated for each individual at each different mean coverage using the formula from Depristo et al. [9]. For running *lcMLkin*, the underlying allele frequencies at each locus were estimated from CEU 1000 Genomes Phase 1 genotype calls [1].

#### 3.2.3 Genomes Phase 3

We obtained previously estimated genotype likelihoods for 2, 535 individuals from 27 different populations sequenced as part of the 1000 Genomes project Phase 3 (http://ftp.1000-genomes.ebi.ac.uk/vol1/ftp/release/20130502/supporting/genotypelikelihoods/shapeit2/). This includes a number of individuals who are known to be related as inferred from previous SNP array genotyping. In general, sequence coverage for this data is likely to be low, though exact mean coverage values were still being compiled during this study.

We sub-sampled 13 putatively non-admixed populations for which there are 48 known pairs of related individuals: Dai (CDX), Southern Han (CHS), Esan (ESN), British (GBR), Gujarati (GIH), Gambian (GWD), Indian (ITU), Kinh (KHV), Luhya (LWK), Mende (MSL), Punjabi (PJL), Tuscan (TSI), Yoruba (YRI). *lcMLkin* was applied to each population separately. Allele frequencies were estimated by applying the Bayesian algorithm described by Depristo et al.[9] and counting the number of variant alleles for the combination of genotypes in the population with the highest posterior probability. Note that there are three pairs of related individuals that are not described in the 1000 Genomes pedigree files [NA19331/NA19334 sibling-sibling in LWK, NA20882/NA20900 parent-offspring in GIH, NA20891/NA20900 parent-offspring in GIH] but have been found elsewhere (http://bloggoldenhelix.com/bchristensen/svs-population-genetics-and-1000-genomes-phase-3) and are confirmed in our study.

In addition to inferring relatedness with *lcMLkin*, it was also inferred for each population with PLINK [33], which was given either the highest likelihood (best) genotypes from single-sample calling, or genotypes obtained through multisample calling that had been conducted as part of the 1000 Genomes project.

## 4 Results

### 4.1 Simulated Pedigrees

We first tested our approach to infer *K*, *ϕ* and *r* under different genome coverage conditions using simulated data consisting of 100,000 independent loci for pedigrees with founders from a non-admixed, non-inbred population, where mean genome coverage ranged 2-20x. When utilizing only the most likely genotype, the estimated 2*ϕ* = *r* is approximately half the true value when mean coverage is 2x in both pairs of samples, and is still slightly underestimated even at 10x (Figure 2). Only at ∼ 20x is *r* correctly estimated. However, when summing over all possible genotypes using *lcMLkin* our estimates of *r* are essentially unbiased even when at 2x, and it appears to be possible to discriminate between 5^th^ degree relatives and unrelated pairs of individuals using this number of loci.

**Figure 2:**
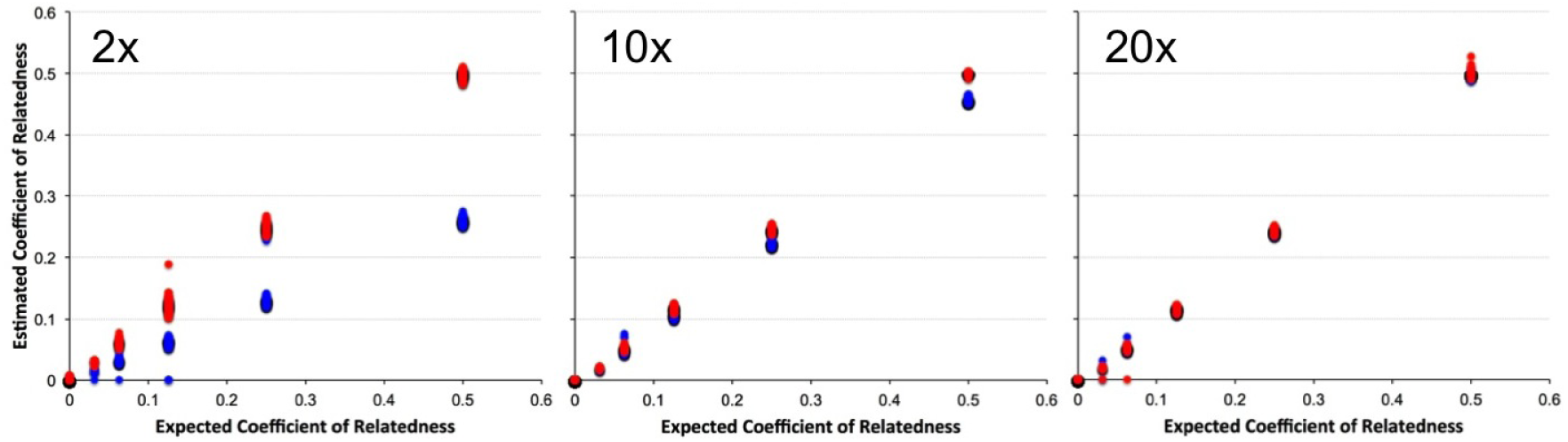
Coefficient of relatedness, *r*, estimated by our method from simulated 2,10 and 20X coverage data versus the known *r*. Blue dots are estimates using only the genotype with the highest likelihood, and red dots are estimates from summing over all possible genotypes in *lcMLkin*

In addition, when we look not only at the estimate of 2*ϕ* = *r* but also *K* (via *k*_0_), we see that the approach of *lcMLkin* clearly distinguishes between sibling-sibling and parent-offspring relationships at 2x coverage, while using only the best genotype results in confounding estimates of *k*_0_ (Figure 3).

**Figure 3:**
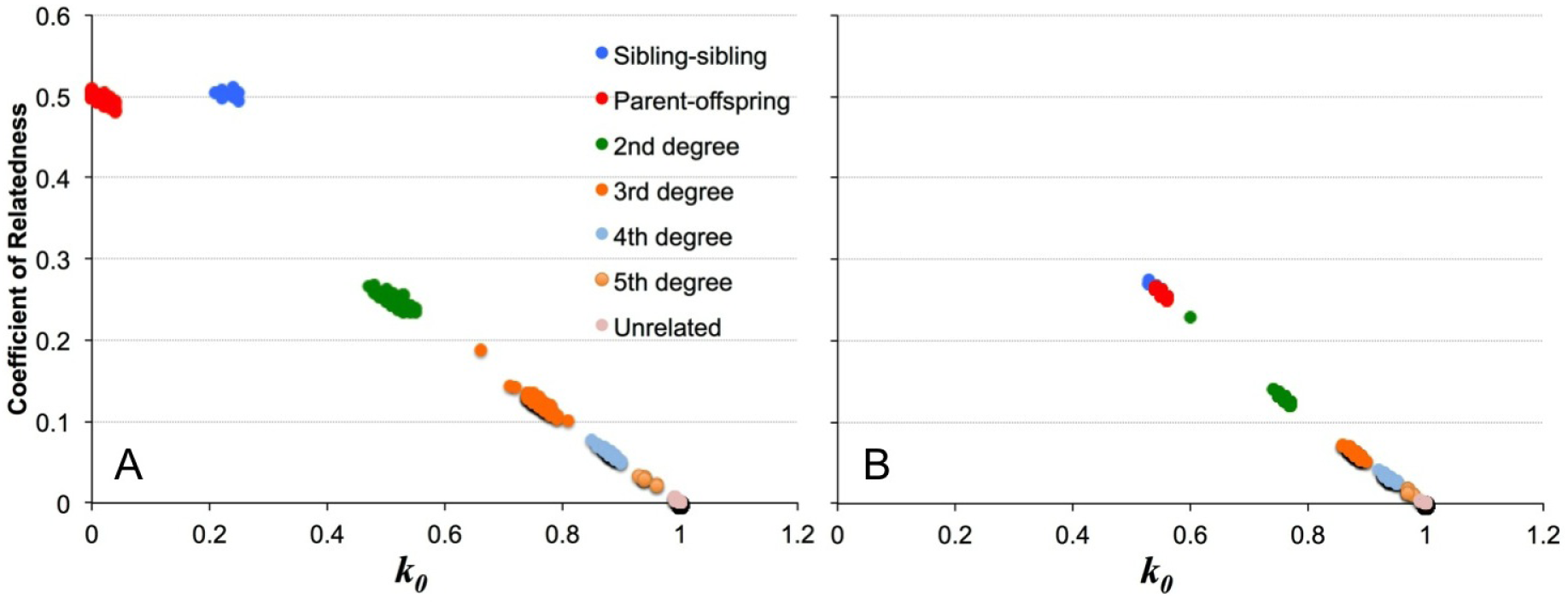
*r* versus kinship coefficient *k*_0_ estimated from simulated 2x coverage data using the sum over all genotypes (A) and just the best genotypes (B). Blue=full siblings, red=parent-offspring, green=2^nd^ degree, orange=3^rd^ degree, sky blue=3^rd^ degree, pale orange=5^th^ degree, pink=unrelated

### 4.2 CEPH pedigree 1463

In order to examine how *lcMLkin* would perform with a more realistic error structure (including mapping error) we examined five individuals from CEPH pedigree 1463 for which high coverage (∼50x) 2^nd^ generation sequencing data has already been generated, and down-sampled sequence reads from each individual at 10, 000 independent SNPs to various mean coverage values ranging from 2-20x. Population allele frequencies were estimated from CEU 1000 Genomes Phase 1 data.

Figures 4 and 5 show a similar pattern to the simulated data described above, with using the best genotype resulting in an underestimate of 2*ϕ* = *r* and an inability to distinguish parent-offspring and sibling-sibling relationships with low coverage, while summing over all genotype likelihoods using *lcMLkin* results in largely unbiased estimates regardless of coverage, indicating our method works well, even when the structure of errors is potentially more complex than what is represented in our simulated data.

**Figure 4:**
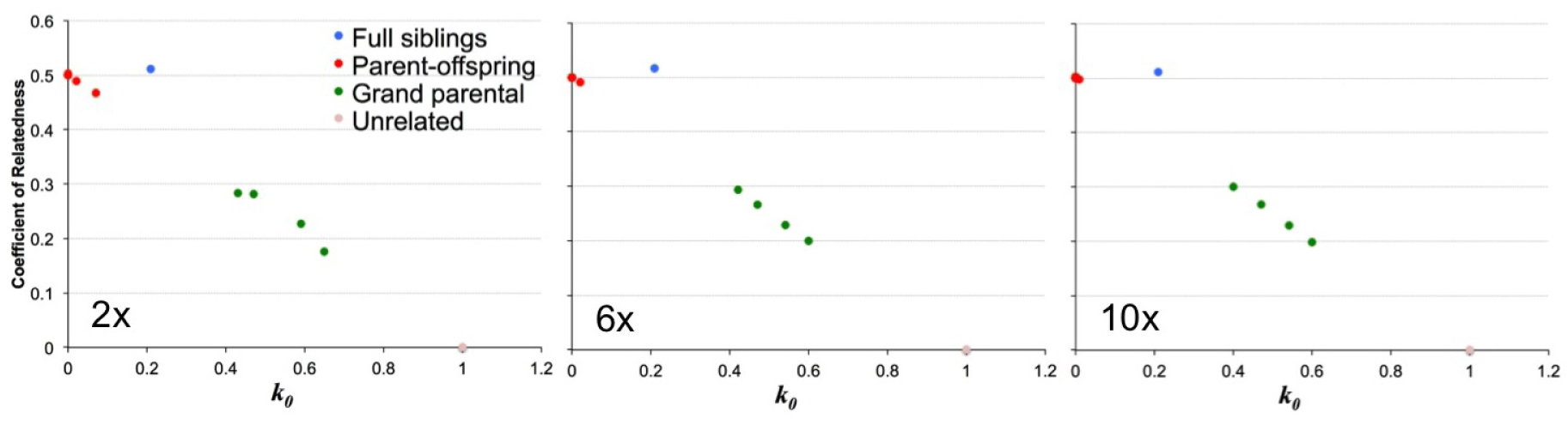
*r* versus kinship coefficient *k*_0_ estimated for pairs of CEPH pedigree 1463 individuals down-sampled to 2x,6x, and 10x mean coverage using the most likely genotype at each SNP. Blue=full siblings, red=parent-offspring, green=grand parental, pink=unrelated

**Figure 5:**
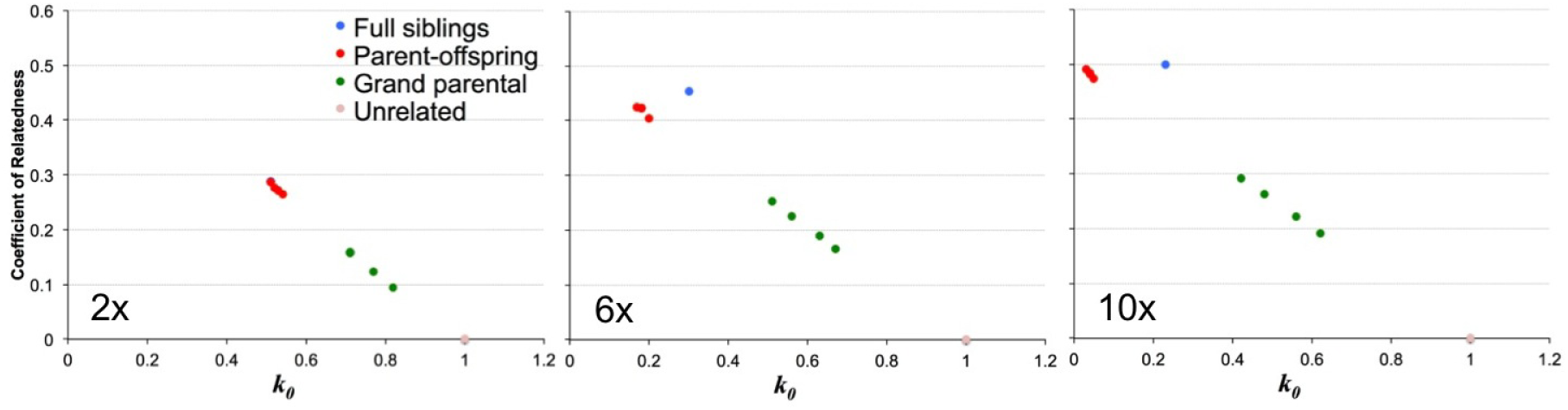
*r* versus kinship coefficient *k*_0_ estimated for pairs of CEPH pedigree 1463 individuals down-sampled to 2x,6x, and 10x mean coverage summing over all possible genotypes at each SNP. Blue=full siblings, red=parent-offspring, green=grand parental, pink=unrelated

In order to examine how incorrect allele frequencies may affect inference by *lcMLkin*, we used the Balding-Nichols model [5] to perturb the population allele frequencies at each SNP with *F*_*ST*_ = 0.01, 0.05 and 0.1. We then re-ran our analysis for the data down-sampled to 2x coverage (Figure 6). For *F*_*ST*_ = 0.01 the estimates of 2*ϕ* = *r* and *k*_0_ are still close to the expected value. Increasing *F*_*ST*_ to 0.05 and then 0.1 results in an increasing overestimation of 2*ϕ* = *r* and underestimation of *k*_0_, though interestingly it seems that it would still be possible to identify parent-offspring and sibling-sibling relationships at 2x coverage even when using populations allele frequencies that are highly diverged from the true values.

**Figure 6:**
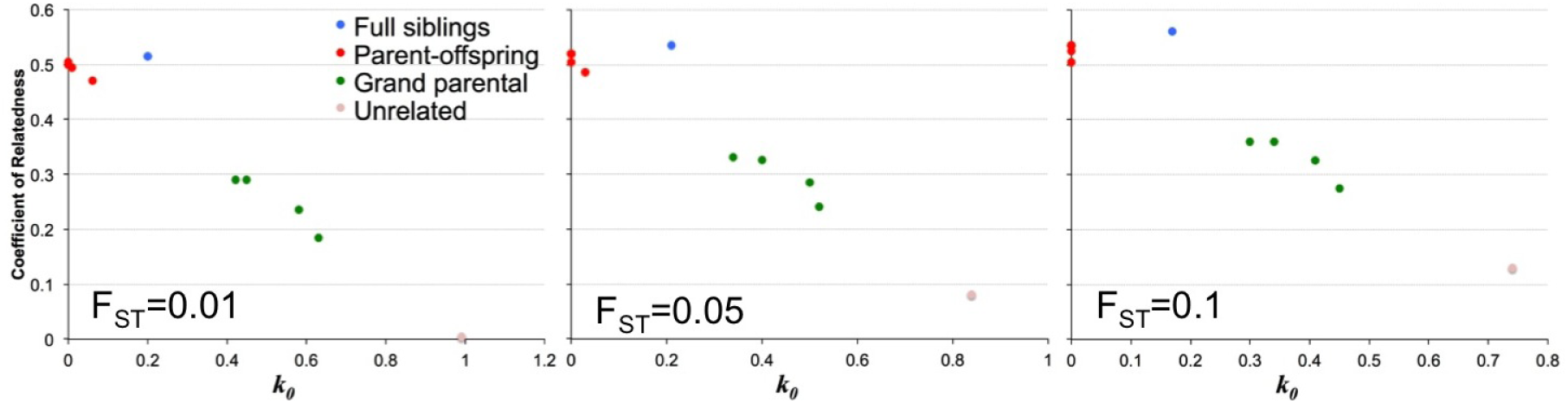
*r* versus kinship coefficient *k*_0_ estimated for pairs of CEPH pedigree 1463 individuals down-sampled to 2x coverage summing over all possible genotypes at each SNP, with underlying population allele frequencies peturbed with *F*_*ST*_ = 0.01, 0.05 and 0.1. Blue=full siblings, red=parent-offspring, green=grand parental, pink=unrelated

### 4.3 1000 Genomes Data

As a final examination of the performance of *lcMLkin*, we analyzed sequence data generated as part of the 1000 Genomes Phase 3 dataset. This dataset contains low coverage sequence data (though the exact coverage for each sample was still being calculated during the writing of this paper) from 48 pairs individuals across 13 putatively non-admixed populations for which there is a know degree of biological relatedness ranging from first to third degree. We applied *lcMLkin* using previously inferred genotype likelihoods to all pairs of individuals within each of the 13 populations at 100,000 independent SNPs. We also applied PLINK [33], a commonly used method of moments estimator to a) the genotype with the highest likelihood for each individual at each SNP and b) the genotype inferred by multisample calling employed by the 1000 Genomes consortium (Figure 7).

**Figure 7:**
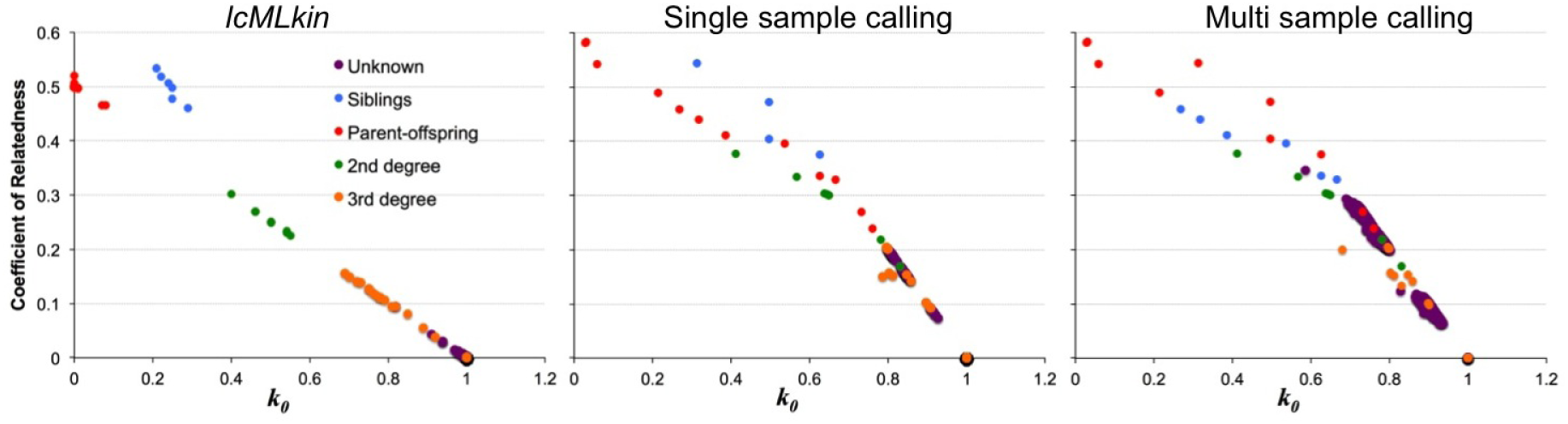
*r* versus kinship coefficient *k*_0_ estimated using *lcMLkin*, PLINK using the genotype with the highest likelihood and PLINK using the genotype inferred by multisample calling for pairs of individuals of known biological relatedness as well as 1000 random individuals of unknown biological relatedness from the 1000 Genomes Phase 3 Project. Blue=full siblings, red=parent-offspring, green=2^nd^ degree, orange=3^rd^ degree, purple=unknown

*lcMLkin* is able to recover all known relationships down to the 2^nd^ degree and most 3^rd^ degree relationships (though a few are estimated to be more unrelated than expected (i.e. lower 2*ϕ*= *r* values)). Pairs of individuals of unknown relationship also generally cluster such that they are inferred to be unrelated (as there are many such pairwise comparisons, only 1,000 random unknown pairwise relationships are plotted in Figure 7 for easy visualization). However, PLINK produces highly inconsistent results, both with single sample and multisample calling, often underestimating 2*ϕ* = *r* and overestimating *k*_0_ for known first to third degree relatives, while a large number of known pairs are inferred to have 2*ϕ* = *r* values indicating they are close to 2^nd^ degree relatives.

## 5 Discussion

We have demonstrated that it is possible to make accurate inference of biological relatedness down to at least three degrees of genealogical separation from 2^nd^ generation sequencing data even when mean coverage is as low as 2x. While our simulations reflect a relatively simple error model, the performance of *lcMLkin* on real data, both under controlled (CEPH pedigree 1463 data) and uncontrolled (1000 Genomes Phase 3 data) conditions, is similar. While being more computationally expensive, *lcMLkin* also vastly outperforms existing methods-of-moment estimators such as PLINK [33]. Further, an efficient and highly-parallel implementation of the EM procedure makes it feasible to apply *lcMLkin*, even to relatively large datasets.

Our approach utilizes information about all possible genotype likelihoods at independent SNPs, rather than assuming a single true genotype. Currently, it is common practice to perform some form of Bayesian multisample calling for large 2^nd^ generation sequencing datasets to infer genotypes [9]. Such approaches inherently assume that each allele sampled from the dataset is randomly drawn from the population. However, this will not be true if related individuals are present. Therefore, when a low-to-medium coverage 2^nd^generation sequencing dataset is collected for which some form of disease variant discovery or population genetic analysis is to be performed, it may be preferable to apply *lcMLkin* to identify such relationships before calling variants.

When large numbers of samples are available from a population of interest, the estimation of population allele frequencies should be fairly robust even with low coverage. Encouragingly, it still appears to be possible to infer first and second degree relatives (data was not available to test further degrees of relatedness) even when the assumed population allele frequencies were highly divergent from the true frequencies. We found no noticeable effect in the estimation of 2*ϕ* = *r* when using allele frequencies from a population that experiences genetic drift with an *F*_*ST*_ of 0.01 from the true frequencies. To put this value in context for humans, average *F*_*ST*_ amongst European countries is 0.004 [31] and Indian ethnicities 0.01 [34]. Thus, our approach may be particularly useful when there are only a few samples to be examined and for which the underlying population allele frequencies are uncertain but for which another population may be a close surrogate (for example modern European frequencies could be used for DNA collected from ancient European specimens). Only with larger *F*_*ST*_ values of 0.1 do estimates of 2*ϕ* = *r* start to show serious biases (though first and second degree relationships still appear as distinct from other possible relationships). At least within humans, such an *F*_*ST*_ would be the equivalent to using allele frequencies from populations of African origin for individuals that are actually of European or Asian origin, and thus is at the extreme end of human population divergence [7].

We also note that as well as only requiring low coverage data, inference appears to also be possible with a relatively modest number of targeted SNPs. Though the variances for estimates for 2*ϕ*= *r* and *K* are higher, we found that *lcMLkin* could distinguish first to third degree relatives from unrelated individuals in simulated data with as little as 1000 SNPs (data not shown). Thus, our approach may be useful for researchers that utilize methods that target smaller amounts of sequence data, such as RAD tag sequencing [3].

While our approach appears to be effective for many realistic situations, there are two situations that may cause biases. If the individuals being examined have ancestry from multiple source populations (i.e. are admixed) this may lead to unrelated pairs of individuals with 2*ϕ*= *r* that are significantly larger than the expected value of 0 (i.e. incorrectly inferred to be related to some degree) [39]. Moltke and Albrechtsen [27] recently described a likelihood-based approach for accounting for such admixed individuals. A natural extension, therefore, would be to extend *lcMLkin* to incorporate this model. However, this will require the exploration of a large number of parameters, which may reduce power to accurately infer 2*ϕ*=*r* and *K* in lower coverage data, especially when individuals are highly (*∼*50%) admixed.

A second situation that may result in incorrect inference of 2*ϕ*=*r* would be populations or specific target individuals that are inbred. Extending the number of *k* coefficients to account for inbreeding [26, 42] could provide more realistic estimates of 2*ϕ*= *r*. However, as with the case of incorporating admixture, this will again increase the parameterization of the model, which may reduce power. In such cases, it may be possible, with some extra computational effort, to provide e.g. credible intervals for parameter estimates by adopting a Gibbs sampling approach in lieu of the existing EM algorithm. This would, at least, allow quantification of uncertainty in the parameters being estimated.

The challenge going forward, therefore, will be to increase statistical power through better resolution of IBD by incorporating information about SNPs in linkage disequilibrium (for example by identifing IBD blocks and thus the distribution of IBD tract length) while accounting for genotyping uncertainty. This would require not only accounting for genotype likelihoods, but also the likelihoods of the haplotypes made up of the individual alleles. Whether this is achievable will determine whether the general approach described here for *lcMLkin* could be extended to allow the inference of more complex biological relationships using low coverage 2^nd^ generation sequencing data.

## 6 Acknowledgments

We thank Gil McVean and Richard Durbin for permission to publish results using 1000 Genomes Phase 3 data. This work is supported by NSF award number 1450606.

